# Multicenter Analysis of Dosing Protocols for Sotalol Initiation

**DOI:** 10.1101/531301

**Authors:** Minakshi Biswas, Andrew Levy, Rachel Weber, Khaldoun Tarakji, Mina Chung, Peter A. Noseworthy, Christopher Newton-Cheh, Michael A. Rosenberg

## Abstract

Sotalol is a Vaughan-Williams Class III antiarrhythmic medication that is commonly used in the management of both atrial and ventricular arrhythmias. Like others in this class, sotalol carries a risk of the potentially lethal arrhythmia torsade de pointes due to its effect of prolonging the QT interval on ECG. For this reason, many centers admit patients for telemetry monitoring during the initial 2-3 days of dosing. However, despite its widespread use, little information is available about the dosing protocols used during this initiation process. In this multicenter investigation, we examine the characteristics of various dose protocols in 213 patients who initiated sotalol over a 4-year period. Of these patients, over 90% were able to successfully complete the dosing regimen (i.e., were discharged on the medication). Significant bradycardia, excessive QT prolongation, and ineffectiveness were the main reasons for failed completion. We found that any dose adjustment was one of the strongest univariate predictors of successful initiation (OR 6.6, 95%CI 1.3 – 32.7, p = 0.021), while initial dose, indication, and resting heart rate or QT interval on baseline ECG did not predict successful initiation. Several predictors of any dose adjustment were identified, and included diabetes, hypertension, presence of pacemaker, heart failure diagnosis, and depressed LV ejection fraction. Using marginal structural models (i.e., inverse probability weighting based on probability of a dose adjustment), we verified that these factors also predicted successful initiation via preventing any dose adjustment, and suggests that consideration of these factors may result in higher likelihood of successful initiation in future investigations. In conclusion, we found that the majority of patients admitted for sotalol initiation are successfully discharged on the medication, often without a single adjustment in the dose. Our findings suggest that several factors predicting lack of dose adjustment could be used clinically to identify patients who could potentially undergo outpatient initiation, although prospective studies are needed to verify this approach.

## Background

Sotalol is a Vaughan-Williams (VW) Class III antiarrhythmic agent that is used in the treatment of atrial^1-3^ and ventricular arrhythmias^4-8^. Composed of a racemic mix of two isomers, the *d* and *l* isomers of sotalol work together to prolong repolarization (*d- and l-*isomer) along with non-selective beta-blocking properties (*l-*isomer). This property is significant as use of isolated d-Sotalol was found to be associated with increased mortality in patients with heart failure in the SWORD study^9^, while use of the racemic mixture is generally considered to be safer in this population. Like other VW Class III agents, sotalol displays reverse use dependence properties, in which heart rate and the QT interval are inversely related such that at lower heart rates, the QT interval prolongs^10^. At doses between 160 and 640 mg/day, sotalol increases QT interval by 40 to 100 ms^11^. In a cohort of 541 patients obtained from electronic medical record review starting sotalol, Weeke el al. found that the average change in corrected QT interval (Bazett formula) was highly variable (3±42 ms at two hours and 11±37ms at 48 hours) following the initial dose^12^.

Due to these properties of sotalol, there is a 1-4% incidence of developing adverse QT prolongation and subsequent torsade de pointes (TdP) in a dose-dependent fashion^13-21^, with risk factors including dose above 320 mg/day, serum creatinine over 1.4mg/dL in women and over 1.6mg/dL in men, history of sustained VT/VF, history of heart failure or coronary artery disease, and female gender placing people at higher risk^14^. As a result, most, but not all, institutions require admission to a monitored telemetry unit for initiation and dose adjustments of sotalol. However, despite multiple observational studies examining patterns of sotalol initiation^12, 14, 22, 23^, no single algorithm exists to guide initiation of sotalol therapy. In this investigation, we examine patterns of sotalol initiation across multiple institutions nationwide to identify predictors of successful initiation.

## Methods

### Population

The National Torsade de Pointes consortium is composed of representatives from the Massachusetts General Hospital, University of Colorado Hospital, Cleveland Clinic, Beth Israel Deaconess Medical Center, and the Mayo Clinic. Patients who were admitted to the hospital for in-patient telemetry monitoring for sotalol initiation for non-research purposes were enrolled at these centers over the period from June 8, 2014 and September 18, 2018. Patients undergoing admission for dose adjustments were excluded. Clinical information was collected on patients over the duration of the hospitalization, including electrocardiography, lab values, and clinical events. Data was collected in a RedCap database; Massachusetts General Hospital served as the data-coordinating center for this investigation. This study is registered with the U.S. National Library of Medicine ClinicalTrials.gov (NCT02439658). The study was approved by institutional review boards (IRB) at all participating institutions.

### Study parameters

Baseline data were collected prior to initiation of sotalol, and included age, sex, demographic information, race, ethnicity, height, weight, body mass index (BMI), past medical history of atrial fibrillation, ventricular tachycardia, essential hypertension, coronary artery disease, congestive heart failure, and presence of a pacemaker or implantable cardioverter-defibrillator (ICD), routine lab values (serum potassium, magnesium, and creatinine), assessment of left ventricular function (i.e., ejection fraction) based on transthoracic echocardiography, nuclear imaging, or cardiac magnetic resonance imaging, and standard electrocardiographic (ECG) rhythm and measurements (heart rate, PR, QRS, QT intervals). ECGs were over-read for accuracy by a cardiologist member of the research team. Underlying atrial rhythm was categorized as atrial fibrillation, flutter, or tachycardia (AF); sinus or atrial-paced (SR) rhythm; or unknown (UNK) if both sinus and atrial arrhythmias were present, or if junctional or ventricular-paced rhythm was present. An ECG was also categorized as UNK if greater than 3 atrial or ventricular premature complexes were noted. UNK ECGs were excluded from ECG analysis (intervals and rhythm), although other clinical information for these patients was included. The QT was corrected using Fridericia^24^ formula (QT/(RR interval)^1/3^. All patients were included regardless of existence of prior bundle-branch block or ventricular pacing, with QRS duration examined as a covariate. Data were also collected regarding timing of electrical or chemical (spontaneous) cardioversion.

### Outcome

We considered a three-day course of 5-6 doses of sotalol to be successful completion of the protocol. Patients who were discharged on sotalol after shorter courses of initiation were excluded; otherwise, any patient who stopped the initiation protocol prior to the 5^th^ or 6^th^ dose and/or was not discharged on sotalol was considered a protocol failure. Reasons for failure were categorized by study investigators into ‘ineffective’, ‘bradycardia’, ‘QT prolongation’, ‘other intolerance/side effect’, and ‘other/unknown’, although the decision about continuation or cessation of sotalol was made by the treating physicians, independently of the research study.

### Analysis

All analyses were conducted on a single dataset downloaded from the central RedCap database on September 20, 2018. Univariate analysis was conducted using logistic regression with successful discharge on medication as the outcome to identify individual predictors of successful initiation. Due to the limited number of unsuccessful initiation protocols (underpowered), multivariate analysis was not performed with successful initiation as the outcome. To examine causal pathway for factors leading to dose adjustment, we used a marginal structural model^25, 26^ based on inverse probability weighting for dose adjustment. First, stepwise logistic regression was performed by dose across all predictors in order to identify those predicting a single dose adjustment at significance level of p < 0.1. Then, generalized estimating equations (GEE) were applied using a logit link and subject-level clustering across doses, to examine the inverse of the predicted probability of a dose adjustment by dose for each subject on probability of successful initiation of sotalol. All statistical analysis was performed using Stata IC (Version 15.1, StataCorp, LLC, College Station, TX).

## Results

Between June 8, 2014 and September 18, 2018, 213 subjects were admitted to five participating medical centers (range 1 – 100 per center). Atrial fibrillation (AF) was the most common indication, and most patients were in AF at the start of the initiation protocol (Table 1). Sotalol was initiated in more men than women, although there were no statistically significant differences in successful completion of the loading protocol between sexes.

**Table 1:** Patients Characteristics

|  |  |
| --- | --- |
|  | Total (N = 213) |
| Age (years) | 66.2 ± 10.5 |
| Ethnicity |  |
| - Hispanic/Latino | 8 (3.8%) |
| - Non Hispanic/Latino | 201 (94.4%) |
| - Unknown | 1 (0.5%) |
| Race |  |
| - Caucasian | 206 (96.7%) |
| - African-American or Black | 3 (1.4%) |
| - Asian | 1 (0.5%) |
| - More than one race | 3 (1.4%) |
| Medications on Admission |  |
| - Beta Blocker | 110 (51.6%) |
| - Calcium Channel Blocker | 33 (15.5%) |
| - Diuretic | 49 (23.0%) |
| Medical History |  |
| - Atrial fibrillation | 185 (86.9%) |
| - Ventricular tachycardia | 33 (15.5%) |
| - CHF | 34 (16.0%) |
| - HTN | 112 (52.6%) |
| - Type II DM | 19 (8.9%) |
| - CAD | 50 (23.5%) |
| - Pacemaker | 19 (8.9%) |
| - Implantable cardioverter-defibrillator | 26 (12.2%) |
| Indication for Sotalol Loading: Atrial fibrillation | 181 (87.4%) |
| Most recent LV EF assessment |  |
| - LV EF > 35% | 159 (88.8%) |
| - LV EF ≤ 35% | 20 (11.2%) |
| Previously on sotalol? (Y/N) | 16 (7.6%) |
| Admission ECG |  |
| - Sinus Rhythm or atrial paced | 99 (48.5%) |
| - HR | 84.6 ± 23.7 |
| - PR | 184.6 ± 44.4 |
| - QRS | 105.8 ± 28.5 |
| - QTc (Fridericia) | 466.2 ± 50.9 |
| Pertinent Laboratory Data |  |
| - Potassium | 4.3 ± 0.4 |
| - Magnesium | 2.0 ± 0.2 |
| - Creatinine Clearance | 0.97 ± 0.26 |
| Sotalol Loading |  |
| - Starting dose |  |
| - 40mg | 8 (3.8%) |
| - 80mg | 134 (62.9%) |
| - 120mg | 69 (32.4%) |
| - 160mg | 2 (0.9%) |
| - Median Number of doses received (IQR) | 5 (5-6) |
| - No dose adjustment (after 5-6 doses) | 113 (53.1%) |
| - Cardioverted electrically | 88 (41.3%) |
| - Cardioverted chemically | 27 (12.7%) |
| - Discharged on Sotalol | 20 (9.4%) |

113 patients (53.1%) completed a 5 or 6 day initiation protocol without any adjustment in the dose—40mg: 1 (0.9%), 80mg: 64 (56.6%), 120mg: 47 (41.6%), 160mg: 1 (0.9%). 66 patients (31.0%) had at least one adjustment over the 5 or 6 dose protocol, and 34 patients (16.0%) stopped the protocol before reaching the 5^th^ or 6^th^ dose. Figure 1 shows the various dosing trajectories for patients who had at least one change, and average numbers by dose. There was no obvious other pattern than no adjustment through the population, although the most common doses were 80mg and 120mg, and most adjustments were increasing or decreasing between these values. 20 patients (9.4%) were not discharged home on the medication. Of these, 11 had stopped prior to the 5^th^ dose, 2 had no dose adjustment and stopped after the 5^th^ dose, and 7 had at least one dose adjustment and stopped after the 5^th^ dose. Of the reasons for unsuccessful initiation, the most common were ‘ineffective’ (8 patients), ‘QT prolongation’ (7 patients), ‘bradycardia’ (4 patients), and ‘other/unknown’ in 1 patient.

**Figure 1.**
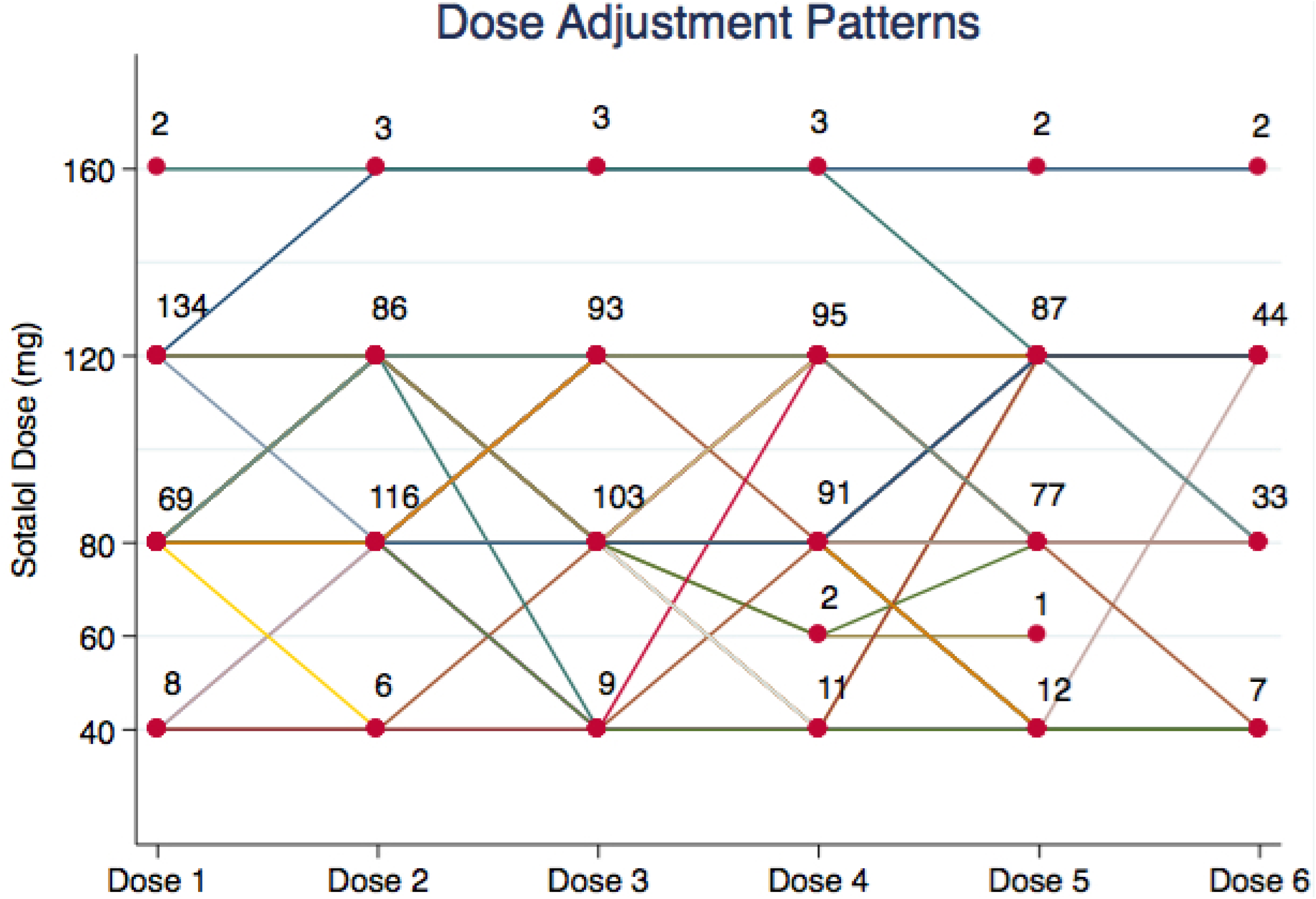
Dose adjustment patterns for sotalol. Numbers indicate patients receiving that dose at each time point. Not shown are patients who had no dose adjustment during the hospitalization.

105 (51.5%) of patients were in atrial fibrillation or flutter on initiation of sotalol, and of these, 61 (66.3%) underwent electrical cardioversion and 20 (19.1%) spontaneously converted to sinus rhythm. There was no significant effect of electrical or spontaneous conversion to sinus on the probability of discharge on sotalol.

In univariate logistic regression, patients with no adjustment in the dose were more likely to be discharged on sotalol than those who had at least one dose change (OR 6.6, 95%CI 1.3 – 32.7, p = 0.021), and patients with a higher starting dose had a higher probability of discharge [120mg: 94.2% (CI 88.7 – 99.7%), 80mg: 89% (CI 84.4-94.7%), 40mg: 75% (CI 45.0-1.05%)], although the latter difference did not reach statistical significance (p = 0.22). Of the predictors examined (Table 2) lack of any dose adjustment was among the most predictive of successful initiation. History of hypertension and use of a separate beta blocker were also predictive of successful discharge on sotalol, although it is possible that the mechanism of these factors was by decreasing the dose changes of sotalol (see below).

**Table 2.** Prediction of successful initiation of sotalol. Based on logistic regression.

|  | **OR** | **CI** | **p value** |
| --- | --- | --- | --- |
| No Dose Adjustment | 6.58 | 1.33 – 32.71 | 0.021 |
| Indication | 0.69 | 0.19 – 2.58 | 0.584 |
| HTN | 0.25 | 0.08 – 0.77 | 0.016 |
| DM | 1.95 | 0.25 – 15.47 | 0.526 |
| PPM | 1.95 | 0.25 – 15.47 | 0.526 |
| CHF | 0.74 | 0.23 – 2.35 | 0.606 |
| Beta blocker | 3.58 | 1.25 – 10.24 | 0.017 |
| Calcium channel blocker | 3.78 | 0.49 – 29.23 | 0.203 |
| LowEF | 0.87 | 0.18 – 4.14 | 0.860 |
| Sinus Rhythm | 1.70 | 0.64 – 4.50 | 0.289 |
| HR | 1.00 | 0.98 – 1.02 | 0.917 |
| QTc (Fridericia) | 1.00 | 0.99 – 1.00 | 0.297 |

Since dose adjustments appeared to play a significant role in the overall successful initiation of sotalol, we examined the role of this decision as a marginal structural model, using inverse probability weighting for dose adjustment. After reshaping the dataset to allow modeling by dose number, we identified that the dose number (OR 0.74, 95%CI 0.64-0.88), presence of hypertension (OR 2.0, 95%CI 1.2 – 3.4), use of calcium channel blocker (OR 2.1, 95%CI 1.2 – 3.8), use of a separate beta blocker (OR 0.63, 95%CI 0.38 – 1.0), and presence of a pacemaker (OR 0.17, 95%CI 0.04 – 0.72) were all predictors for making a dose adjustment at p < 0.1. From the multivariate logistic regression model, we calculated inverse probability of a dose adjustment by dose number, and examined the role in predicting successful initiation of sotalol using generalized estimating equations. This model demonstrated that predictors of dose adjustment were also successful at predicting the probability of successfully completing the initiation protocol of sotalol (OR 0.977, 95%CI 0.960 – 0.995, p = 0.014).

## Discussion

In this multicenter observational study of sotalol initiation protocols, we found that regardless of the dose protocol, the vast majority of patients (> 90%) were able to successfully initiate sotalol during the loading protocol, and that over 50% of the population was able to do so without any adjustments to the medication. We found a large range of variability in the dose adjustments made during initiation of sotalol, but also found that patients in whom dose adjustments were made were less likely to be discharged on the medication. Several factors were associated with less frequent dose adjustments, including presence of a pacemaker, beta blockers or calcium channel blockers, hypertension, and diabetes, and could be examined in future prospective studies aimed at minimizing dose adjustments toward the goal of improving likelihood of sotalol initiation, as well as targeting specific patients for outpatient initiation.

Unlike the related VW class III anti-arrhythmic medication, dofetilide, the initiation protocol for sotalol is poorly defined, and at some institutions, patients being initiated on sotalol are not admitted for any monitoring during the initiation process at all. Several groups examined the safety of in-patient sotalol initiation over 2 decades ago, and noted a higher rate of events than we identified. Maisel and colleagues examined adverse cardiac events in 72 trials of sotalol 80mg twice a day for atrial fibrillation^22^, and found that adverse cardiac events occurred in 18 percent: bradyarrhythmias occurred in eight (11 percent), ventricular arrhythmias in two (2.8 percent), and QT prolongation in one (1.4 percent). The risk was greatest within the first 24 hours of therapy and in patients with a previous myocardial infarction (MI)^22^. Chung et al. also examined the risks of sotalol initiation in 120 patients admitted to a single center^23^, and found that complications occurred in 25 patients (21 percent), which triggered a change in therapy (dose reduction or cessation) in 21. The most common complications were bradycardia in 20 (including a heart rate below 40/min in 13), excessive prolongation of the QT interval in eight, and new or increased ventricular arrhythmias in seven (including two cases of TdP). Complications occurred within the first three days in 22 of the patients (88 percent)^23^. Both studies identified a rate of complications higher than our study, which may reflect greater attention to dose effects in the more contemporary period, although neither study examined dose-by-dose granularity as we did, and thus we cannot determine this effect precisely.

Our finding that over half of patients were able to successfully initiate sotalol without a single dose adjustment has potential implications for future efforts at outpatient sotalol initiation protocols. Noteworthy is that patients with a pacemaker present were significantly less likely to have a dose adjustment, and more likely to successfully initiate sotalol than those without, perhaps due to less bradycardia, which was a key reason for failure of initiation. Given the continued advances in remote monitoring of cardiac implantable devices^27, 28^, as well as development of wearable/portable devices for measuring QT interval^29, 30^, it seems that our findings support the possibility that these patients could undergo outpatient initiation of sotalol, although more work is clearly needed.

The role of dose adjustments in patients being initiated on sotalol should be contrasted with the other VW class III medication for which patients are admitted for initation, dofetilide. For one, patients starting dofetilide are generally started at the highest dose, with adjustments downward based on QT interval^31^. As observed in this study, most patients initiated on sotalol are started at 80mg twice a day, which is the lowest dose at which antiarrhythmic activity (class III) takes place, and either increased or maintained at the same dose over the course of the 3 days of monitoring. Importantly, we found that making any dose adjustments decreased the probability of successful initiation. Of course, whether the decision to make a dose adjustment was made by providers seeking to achieve a pre-determined dose, or whether the decision was made to adjust the dose because of concerns of toxicity or ineffectiveness that ultimately led to discontinuation of the medication could not be determined by our study design. Within the confines of an observational study, such a determination cannot be made definitively, and requires a prospective trial with a predetermined dosing protocol to define the direction of causality. We believe this future investigation would be important, as dose adjustment targeting ineffectiveness rather than toxicity could potentially be made in an outpatient setting over a longer time period, as is done in other non-class III anti-arrhythmics. Such an approach could be much more cost-effective than present standards for in-patient monitoring.

There are several important limitations in this investigation. First, as noted above, because our study was observational in nature, we were unable to draw more definitive conclusions about the role of dose decisions in successful initiation of sotalol. There is an obvious issue of ‘cause-and-effect’ related to making these adjustments, and while certain findings, such as the presence of a pacemaker decreasing the probability of a dose adjustment and increasing probability of successful initiation of sotalol, make intuitive sense; others, such as the associations of calcium-channel blockers and beta blockers with dose adjustments, are more difficult to explain. These associations would seem to be more likely to be spurious, and likely due to other unmeasured effects specific to certain patients, but only through future prospective study can we know for certain. Second, we did not have medium or long-term follow-up, and thus were unable to determine whether patients who did successfully initiate sotalol had long-term success with the medication, free of complications. Such follow-up is of obvious importance for drawing conclusions; however, understanding the factors that impact successful initiation in the short-term is also important. For both patients and providers, the time wasted when a patient is admitted for sotalol initiation and then leaves without being on the medication is a burden both in time and money. Efforts to minimize this outcome are clearly desired, regardless of whether sotalol has long-term success in management of atrial or ventricular arrhythmias. Finally, our use of a multicenter study design had limitations in the depth of information that could be collected about patients. Newer approaches by our group^32^ and others to use machine learning to analyze high-density data^33-35^, such as from telemetry, implanted cardiac devices, and other continuous data streams could provider greater prediction than standard ECG or clinical factors, but are more difficult to collect across institutions and monitoring platforms. Future approaches to include this information could open a treasure trove of data that could provide additional guidance on dose management for medications like sotalol.

In conclusion, we found that the majority of contemporary patients admitted at multiple medical centers for monitoring during initiation of sotalol were successfully discharged on the medication, including over half without a single dose change. We found that among other factors, making dose adjustments was a key risk factor for failed initiation, and that patients with pacemakers already implanted were less likely to have dose adjustments, and more likely to be discharged on sotalol. Further prospective studies are needed to examine the role of dose protocols toward the possibility of safe outpatient sotalol initiation.

## Acknowledgements

This work was support by grants from the NIH NHLBI (MAR: 5K23 HL127296, CNC: R01 HL 143070).

